# Investigation of ice nucleation properties of *Pseudomonas syringae* bacterium and insoluble low molecular weight substances

**DOI:** 10.1101/2023.12.09.570762

**Authors:** D.E. Vorobeva, M.A. Majorina, N.U. Marchenko, B.S. Melnik

**Affiliations:** Institute of Protein Research, Russian Academy of Sciences, 142290 Pushchino, Moscow Region, Russia; Shemyakin–Ovchinnikov Institute of Bioorganic Chemistry, Pushchino Branch, Russian Academy of Sciences, 142290 Pushchino, Moscow Region, Russia

**Keywords:** freezing temperature of water, temperature of coexistence of ice and water, *Pseudomonas syringae*, ice nucleator

## Abstract

Control of the water freezing process is considerable in different fields of science and technology: from the artificial snow production to the cryopreservation of biological materials. To date, there is no conventional theory that predicts the influence of various biological and non-biological ice nucleators on the formation of ice and, accordingly, on the freezing point of supercooled water. In this work, we investigated the influence of bacterium *Pseudomonas syringae*, a biological ice nucleator, and heterodisperse insoluble powders of low molecular weight substances on the process of water freezing. AgCl, ZnO and SnO_2_ were found to be ice nucleators. This property has not been described previously in the literature. It has also been established that insoluble low molecular weight substances affect both the freezing point of water and the temperature of coexistence of water and ice.

## Introduction

Ice formation can occur homogeneously, i.e. in pure water, or heterogeneously, if the water contains substances and surfaces that contribute to formation of the ice nuclei. Homogeneous water freezing occurs at very low temperatures (around -30°C), while heterogeneous freezing of water can occur at temperatures close to zero degrees Celsius (Finkelstein et al., 2022). Despite the fact that many works have been devoted to the study of ice formation at near-zero temperatures (John Morris & Acton, 2013; Ogawa & Osanai, 2020; Roy et al., 2021), there is still no generally accepted theory that could describe this process in detail.

Control of the water freezing process is important in various fields of science and technology, for example, in the production of artificial snow at sports facilities (Baloh et al., 2019), in the design of ice-phobic materials for the aviation industry(Shen et al., 2019), in the cryopreservation of biomaterials (Karnieli, 2016; Murray & Gibson, 2022). Therefore, a lot of experimental work with antifreeze, ice nucleators and substances affecting the process of water freezing can be found in the literature (Bian et al., 2023; John Morris & Acton, 2013; Thiel & Madey, 1987).

Substances facilitating the water freezing at near zero temperatures (ice nucleators) have been known since the 40’s of the 20^th^ century. B. Vonnegut discovered that silver iodide (AgI) promotes the formation of ice nuclei and initiates the process of ice formation at a relatively high sub-zero temperature (from -4°C to -8°C) (Vonnegut, 1947). Later, the ability of various substances to cause ice nucleation was confirmed. Diamond nanoparticles (Han et al., 2008), oxidized carbon nanomaterials (Whale et al., 2015), feldspar particles (Atkinson et al., 2013), as well as biological objects such as proteins (Hansen et al., 2023; Hartmann et al., 2022), steroids (Sosso et al., 2022), tree pollen (Gute & Abbatt, 2020), bacterias (Majorina et al., 2022; Maki et al., 1974) shown to stimulate ice formation.

Comparative studies of various substances are important for understanding the general principles of the nucleators influence on the process of water freezing. In this work, we compared the effect of *Pseudomonas syringae* bacterium and heterodisperse powders of insoluble oxides of transition and post-transition metals (CuO, TiO_2_, Fe_2_O_3_, Al_2_O_3_, SiO_2_, Cr_2_O_3_, ZnO, SnO_2_, Ni_2_O_3_), and silver salts AgI, AgCl on the process of ice formation.

Previously, we designed and tested the device that allows us to study the process of water freezing and ice melting in relatively large volumes of liquid (about one milliliter) (Majorina et al., 2022). The device determines three parameters with an accuracy of several hundredths of a degree: the water temperature at the moment of ice nucleation (water freezing temperature T_f_), the temperature of coexistence of water and ice (T_iw_) and the ice melting temperature (T_m_).

Most methods described in the literature are intended to study only one of these parameters. For example, the methods used to study the freezing of liquids in droplets up to 1 microliter (Budke & Koop, 2015; Roy et al., 2021) allow determining only the water freezing temperature. Techniques that use cryoosmometers only measure the temperature of coexistence of water and ice (Grattoni et al., 2008).

In this work, we investigated the effect of *Pseudomonas syringae*, and heterodisperse powders of low molecular weight substances, on the freezing point of water and the temperature of water and ice coexistence. As a result, we showed AgCl, ZnO and SnO_2_ to be ice nucleators; this property has not been described previously in the literature. It was also unexpected that both the bacterium *Pseudomonas syringae* and some insoluble low-molecular substances affected the temperature of coexistence of ice and water.

## Materials and Methods

### The design and protocol of the experiment

The design of the equipment and the features of the experiments are described in detail in the article (Majorina et al., 2022). The only difference concerns the thermostat. In this work, we used a liquid thermostat (Julabo F25), controlled from a computer and capable of changing the temperature in the measuring cell. The thermostat cyclically repeated cooling from +10°C to -18°C, and then heating from -18°C to +10°C at a rate of 0.24 degrees/min. The temperature in the measuring cell (2 ml tube and sample volume 1 ml) was controlled by thermometer (TERMEX LTA), which absolute error equals 0.05°C and the relative error while measuring does not exceed 0.003 °C. Figure 1 shows the curve based on the results of one measurement cycle. The arrows show sections of the curve for determining the freezing temperature of water (T_f_) and the temperature of coexistence of ice and water (T_iw_). Detailed explanations of the features of such a curve are described in (Majorina et al., 2022). Cooling and heating cycles were repeated from 5 to 15 times. Each subsequent measurement was accompanied by change of both the sample and the test tube.

**Figure 1.**
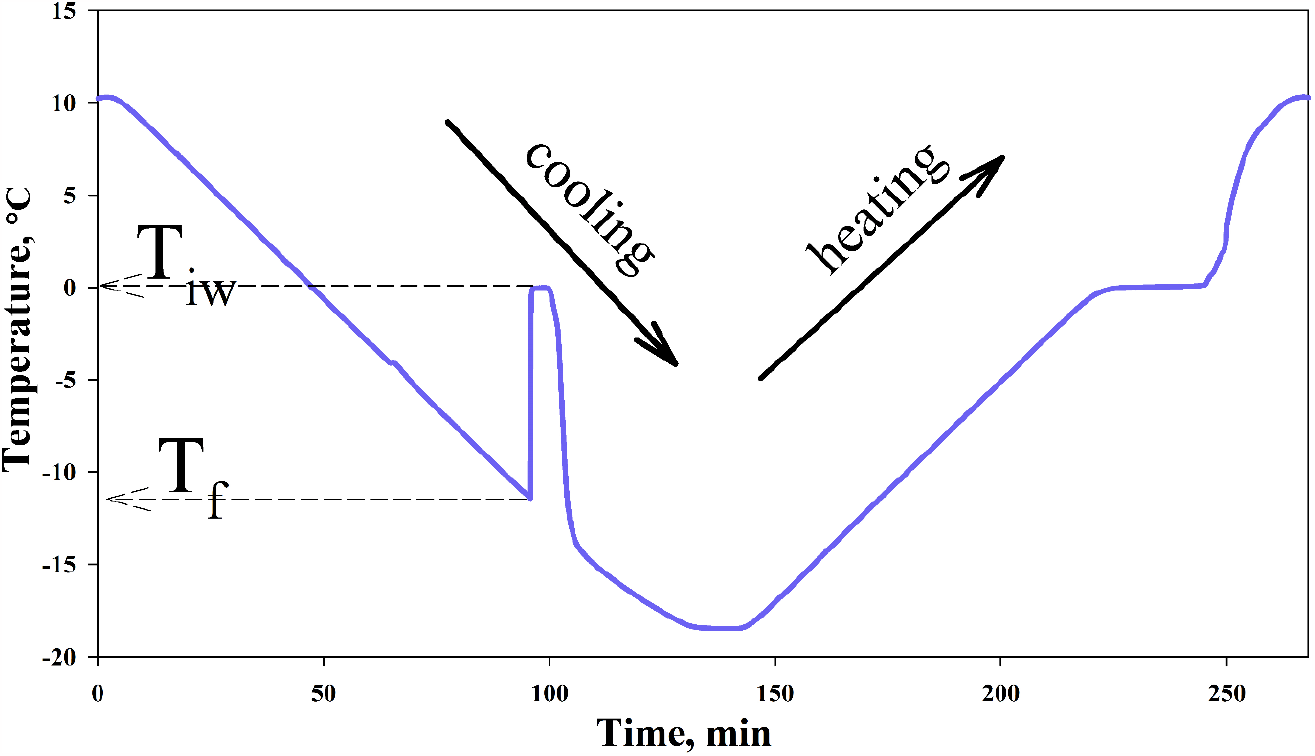
Temperature of a sample in a test tube during a cooling-heating cycle. Cooling from +10°C to -18°C, heating from -18°C to +10°C. The dotted arrow shows how T_f_ – the freezing temperature of supercooled water, and T_iw_ – the temperature of coexistence of water and ice, is detected from the graph.

### Substances used in the work

The study used dry powders of CuO, ZnO, SnO_2_, Al_2_O_3_, SiO_2_, Fe_2_O_3_, TiO_2_, Ni_2_O_3_, Cr_2_O_3_ (Merk, Sigma Aldrich, USA) and AgI, AgCl (Thermo Fisher Scientific, India) with a content of the main component of at least 98%. After dilution the samples with deionized water, the tubes were precipitated on a centrifuge at 3 000 g for sedimentation of substance’s particles.

### Preparation of Pseudomonas syringae culture suspension

*P. syringae* cells (*Pseudomonas syringae* pv. *syringae*) were grown on medium L (yeast extract 5.0 g/l; peptone 15.0 g/l; NaCl 5.0 g/l) at a temperature of 26°C. All the cells were grown in a liquid medium up to cell density of OD_600_ = 1.0 OU, then precipitated on a centrifuge at 6 000 g, and washed twice with a solution of 20 mM Tris-HCl, pH 7.5. The initial cell solution was diluted with a buffer solution of the same composition to the desired concentration. The concentration of cells was controlled by absorption at 600 nm.

## Results

### Research of T_f_ and T_iw_ values of water in the presence of expected ice nucleators

The influence of various substances on the freezing process of water was determined as follows. 1 ml of water was poured into the test tube and cooling-heating cycles were carried out several times (Fig. 1 shows an example of one cycle). Then, the water freezing temperature T_f_ and the coexistence temperature of water and ice T_iw_ were determined. The resulting curve also allows us to determine the melting temperature of ice, but in this research, we did not use these values, since we previously showed that this parameter depends on the design of the equipment and is systematically overestimated (Veselova et al., 2022). Then 2 to 5 milligrams of insoluble substance powder or 50 microliters of *Pseudomonas syringae* suspension were added to the same test tube, and several cooling-heating cycles were also carried out to determine the temperatures T_f_ and T_iw_.

Figure 2 shows the T_f_ of pure water and the T_f_ of water with the addition of *Pseudomonas syringae* culture and AgI, AgCl, CuO, ZnO, SnO_2_, Al_2_O_3_, SiO_2_, Fe_2_O_3_, TiO_2_, Ni_2_O_3_, Cr_2_O_3_ powders. Figure 3 shows the T_iw_ values.

**Figure 2.**
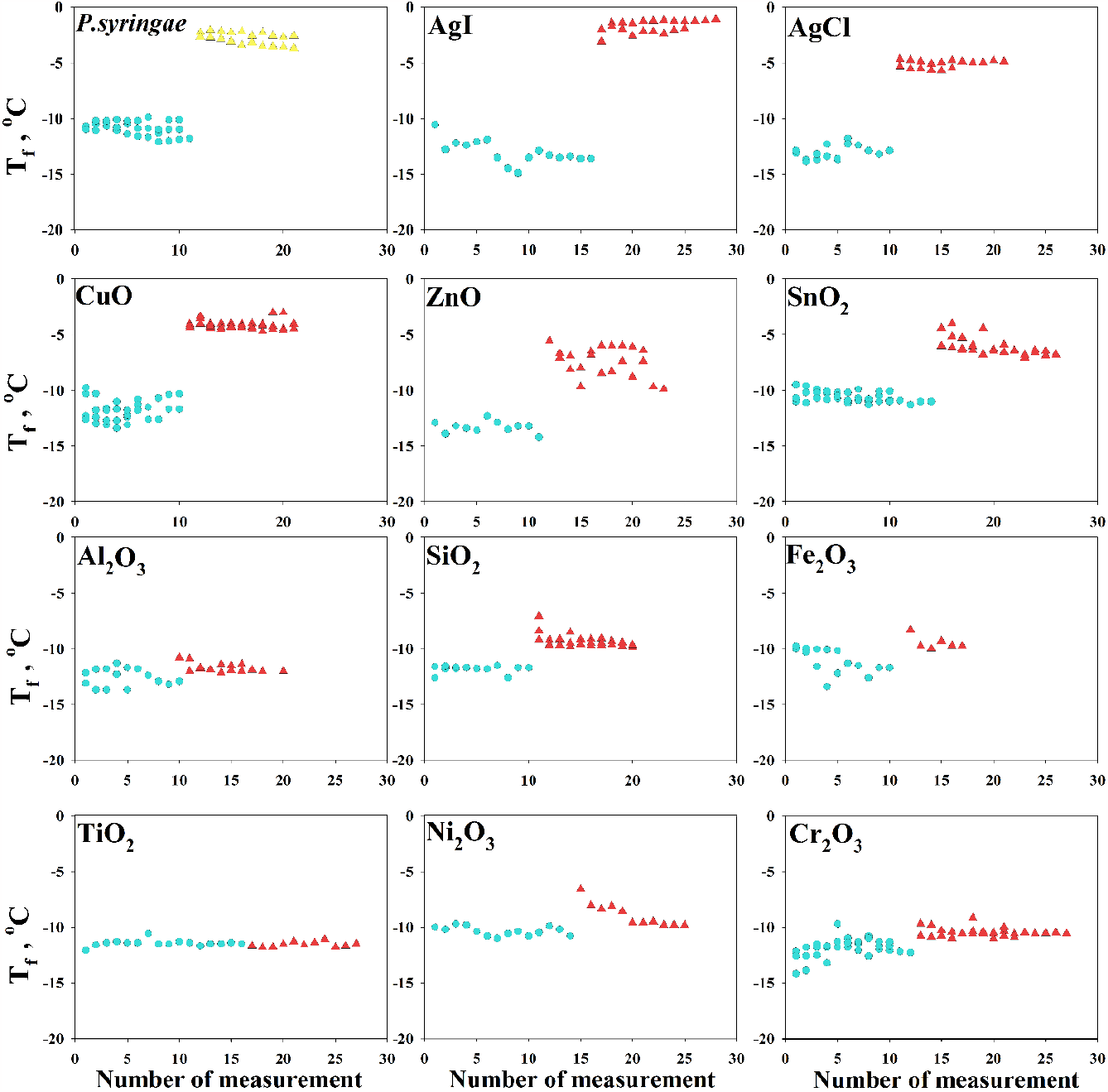
The freezing temperature T_f_ of pure water in a test tube (indicated by circles) and water after adding *Pseudomonas syringae* and AgI, AgCl, CuO, ZnO, SnO_2_, Al_2_O_3_, SiO_2_, Fe_2_O_3_, TiO_2_, Ni_2_O_3_, Cr_2_O_3_ powders (indicated by triangles). Number of measurement corresponds to the number of the cooling-heating cycle.

**Figure 3.**
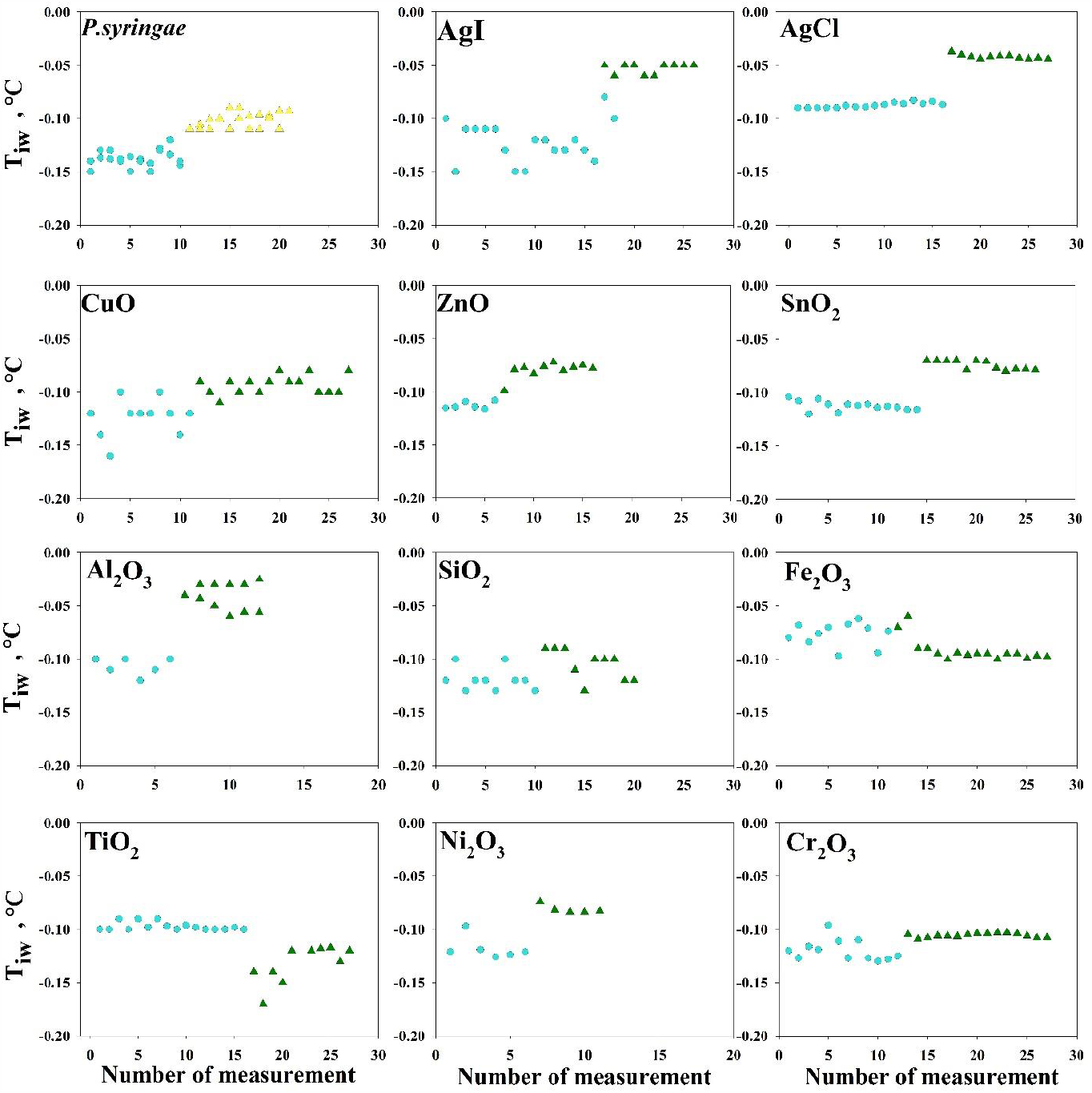
The temperature of water and ice coexistence T_iw_ (indicated by circles) and T_iw_ after the addition of *Pseudomonas syringae* and AgI, AgCl, CuO, ZnO, SnO_2_, Al_2_O_3_, SiO_2_, Fe_2_O_3_, TiO_2_, Ni_2_O_3_, Cr_2_O_3_ powders (indicated by triangles). Number of measurement corresponds to the number of the cooling-heating cycle.

*Pseudomonas syringae* and AgI were known from the literature to be ice nucleators (Curland et al., 2018; Maki et al., 1974). The other test substances were selected from available insoluble metal oxides. Table 1 lists the amounts of added substances and some of their properties specified in the safety data sheets.

**Table 1.** The effect of the studied substances on the freezing parameters of water.

| Substance | Weight, mg<br>(* OU<sub>600</sub> for cells) | T<sub>f</sub>, °C | T<sub>iw</sub>, °C | Solubility in water<br>at 20°C, g/l |
| --- | --- | --- | --- | --- |
| *P. syringae* | 0.01* | -2.390 | -0.097 | soluble |
|  | 0.01* | -3.860 | -0.107 |  |
| CuO | 4.0 | -4.200 | -0.042 | <0.001 (insoluble) |
|  | 4.5 | -4.270 | -0.070 |  |
|  | 3.8 | -5.890 | -0.090 |  |
|  | 4.4 | -5.360 | -0.120 |  |
| Cr<sub>2</sub>O<sub>3</sub> | 4.2 | -10.640 | -0.110 | <0.001 (insoluble) |
|  | 4.5 | -10.320 | -0.100 |  |
| SiO<sub>2</sub> | 4.5 | -9.630 | -0.105 | <0.001 (insoluble) |
| SnO<sub>2</sub> | 3.8 | -4.770 | -0.070 | <0.001 (insoluble) |
|  | 3.9 | -5.350 | -0.070 |  |
|  | 4.2 | -6.330 | -0.075 |  |
|  | 4.0 | -6.240 | -0.110 |  |
| AgI | 4.0 | -1.360 | -0.040 | <0.001 (insoluble) |
|  | 3.8 | -2.150 | -0.053 |  |
| AgCl | 3.6 | -4.897 | -0.042 | 0.00188 |
|  | 4.3 | -5.557 | -0.061 |  |
| ZnO | 4.6 | -7.445 | -0.079 | <0.001 (insoluble) |
| Al<sub>2</sub>O<sub>3</sub> | 4.0 | -11.677 | -0.031 | <0.001 (insoluble) |
| Fe<sub>2</sub>O<sub>3</sub> | 3.9 | -11.944 | -0.092 | <0.001 (insoluble) |
| Ni<sub>2</sub>O<sub>3</sub> | 3.7 | -8.891 | -0.071 | <0.001 (insoluble) |
|  | 4.0 | -6.754 | -0.0814 |  |
| TiO<sub>2</sub> | 4.7 | -11.564 | -0.131 | <0.001 (insoluble) |

T<sub>f</sub> – the freezing temperature of supercooled water, and T<sub>iw</sub> – the temperature of coexistence of water and ice, OU<sub>600</sub> - optical units, the cell solution was measured in a 1cm cuvette at 600 nm. The values of substances solubility in water were taken from branded safety data sheets (SDS) of powders.

Analyzing Figure 2 and the average T_f_ values in Table 1, we can conclude that the addition of *Pseudomonas syringae* and low molecular weight substances AgI, AgCl, CuO, ZnO, SnO_2_, SiO_2_ affected the freezing point of water, increasing it by several degrees. Al_2_O_3_, Fe_2_O_3_, TiO_2_, Ni_2_O_3_, Cr_2_O_3_ did not affect the T_f_ value. To estimate the error of such experiments, we plotted the distribution of water freezing temperatures. Unfortunately, in this plot we could only use data from experiments with pure water, since the number of these experiments is sufficient for analysis (about 300). Figure 4A shows the distribution of T_f_ values for water (marked with circles). By approximating these data with a Gaussian curve, we can conclude that the average freezing temperature of water in a test tube (without the addition of test substances) is -11.4 ± 1.4°C. This method of calculating the freezing point and error is much more accurate than simply calculating the average value. If we assume that after the addition of the test substances, the T_f_ temperatures follow a similar distribution, then ± 1.4°C is the error for all measured T_f_ values. Based on this assumption, we considered that substances affecting the freezing temperature of water by more than 2.8 degrees (AgI, AgCl, CuO, ZnO, SnO_2_, SiO_2_) can be classified as ice nucleators. Al_2_O_3_, Fe_2_O_3_, TiO_2_, Ni_2_O_3_, Cr_2_O_3_ presumably do not cause ice nucleation. The results of adding SiO_2_ to water are ambiguous. The addition of SiO_2_ affected the freezing temperature of water by only 2 degrees (Fig. 2, Table 1) and perhaps this substance is a weak ice nucleator, but taking into account the calculated error of the experiments, it is currently impossible to reliably say whether SiO_2_ is an ice nucleator.

**Figure 4.**
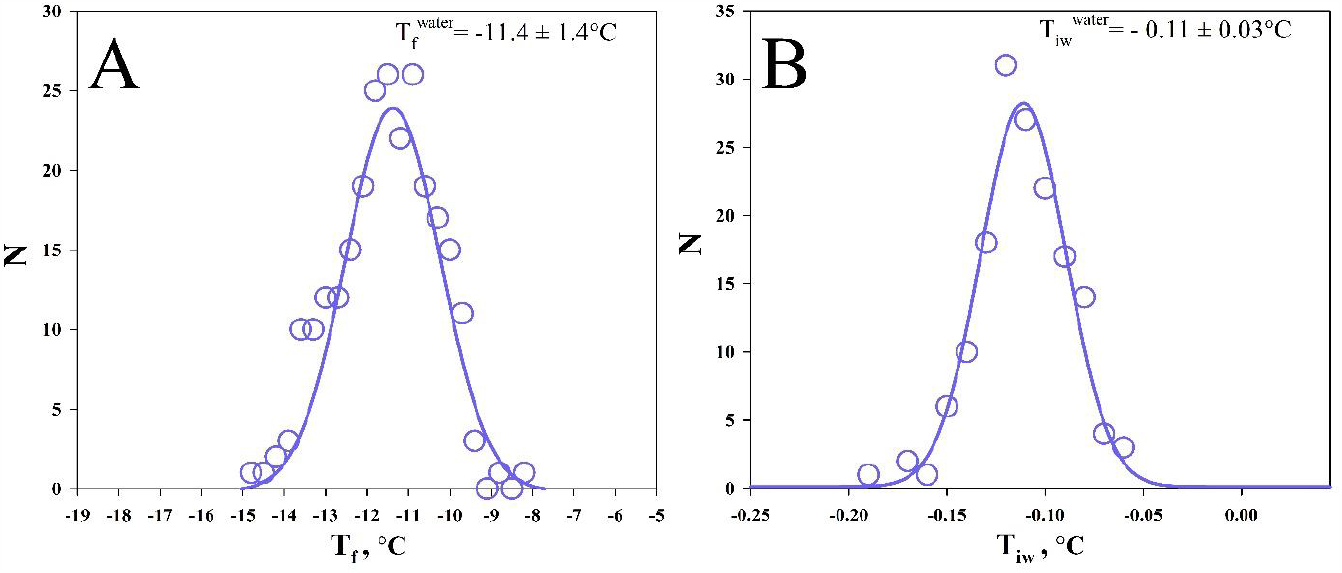
Distribution of the freezing temperature T_f_ (A) and the temperature of coexistence of water and ice T_iw_ (B) values for pure water. N is the number of experiments in which the solution was frozen in the temperature range from T to T+ΔT, ΔT = 0.3°C for T_f_ and ΔT = 0.01°C for T_iw_.

Figure 4B shows the distribution of T_iw_ values, the maximum of the Gaussian curve corresponds to a temperature of -0.11 ± 0.03°C. Figure 3 shows that adding, for example, AgI or Al_2_O_3_ increases the T_iw_ temperature by a value exceeding the error of ± 0.03°C.

### Effect of different amounts of CuO on T_f_ and T_iw_

One of the versions, why insoluble substances in our experiments could affect the T_iw_ temperature is the technical features of the experiment. For example, different amounts of insoluble powder at the bottom of a test tube could have different effects on the heat distribution around the thermometer probe. In the previous article (Veselova et al., 2022) we showed that the distance between the thermometer probe and the bottom or edges of the test tube affects the measurement results. We investigated the effect of different amounts of CuO on the water freezing process. According to the Figure 3, 4 mg of CuO had little or no effect on the T_iw_ value, and therefore this oxide was chosen for the investigation. It can be assumed that an increase in the amount of CuO by several times can affect the T_iw_ temperature more than in Figure 3. Figure 5 shows the dependence of T_f_ and T_iw_ temperatures on different amount of CuO in the test tube. Despite the fact that the amount of CuO in the test tube increased by two orders of magnitude (from 1.5 mg to 170 mg), this did not affect the values either T_f_ or T_iw_. The inset in Figure 5A shows a photo of a test tube containing insoluble CuO powder. It can be seen that the volume of CuO in the test tube varies greatly from 1.5 mg to 170 mg, and if the T_iw_ values depended on the volume of the powder, we would have detected this in experiments. However, different amounts of CuO did not affect T_iw_. It follows that the effect of some test substances on T_iw_ is not explained by technical features of the device or by errors due to different amounts of substance.

**Figure 5.**
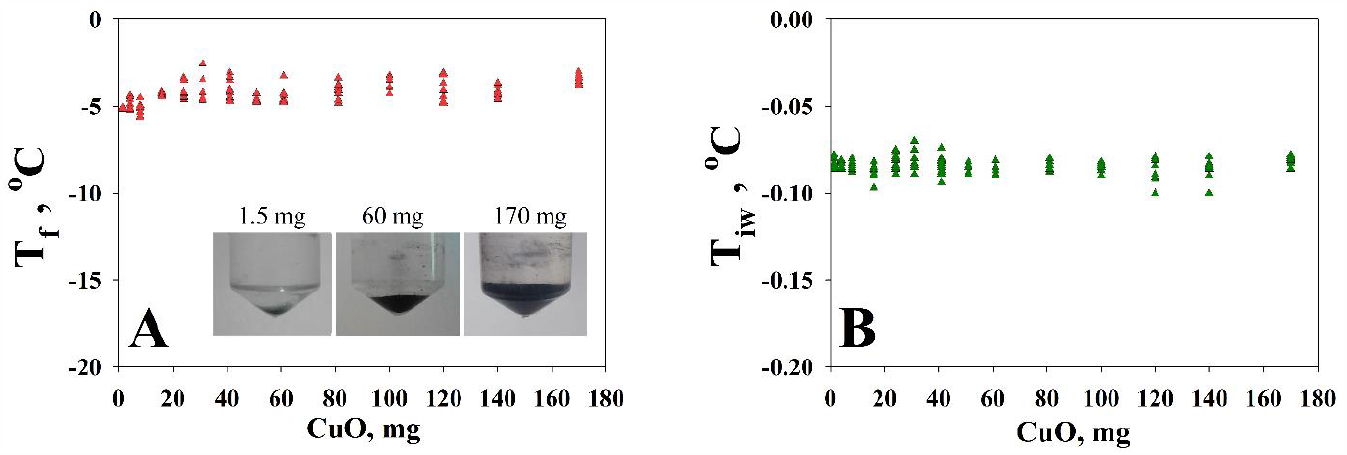
The T_f_ (5A) and T_iw_ (5B) temperature dependence on mass of CuO in the same one test tube. The insertion of figure 5 shows an example of a photo of tubes with water and different amount of sedimented CuO powder (1.5, 60, 170 mg).

## Discussion

As you can see at the Figure 2, conventional nucleators AgI, CuO and *Pseudomonas syringae* distinctly have an influence on the T_f_ temperature. Furthermore, we found AgCl, ZnO, SnO_2_ overt nucleators seeing increasing the temperature of T_f_ due to addition of their powders to the samples. All the studied substances demonstrated different effects on T_f_ either because of their chemical properties or shape of powder particles. Summarizing we can rank founded nucleators by the measure of changes in T_f_ and, thereafter, by the “ice nucleating potential”, from biggest to smallest: AgI, CuO, AgCl, SnO_2_, ZnO.

Results of measuring the temperature of water and ice coexistence T_iw_ turned out to be little bit surprising. Looking at the Figure 3 you can notice nutty feature – after adding some substances, T_iw_ increased, and it did not depend on powder’s ice nucleation activity. For example, Al_2_O_3_ and Ni_2_O_3_ had an effect on T_iw_ unlike the copper oxide CuO. This result is still difficult to explain. It is generally known that solution of any substances in water decreases T_iw_ (for instance, you can recall that any salt melts the ice and coexistence of marine water and ice is detected at temperature lower than 0 °C), and per each mol of solute in water ΔT_iw_ equals 1.86°C (Fink, 2021). We assumed that T_iw_ of sample with insoluble substances will be the same as T_iw_ of pure water, what would confirm the fact of minimal amount of soluble impurity in used powders. We must emphasize that the presence of impurities would have to manifest as decreasing T_iw_ temperature, however, it seems that AgI, AgCl, ZnO, SnO_2_, Al_2_O_3_ and Ni_2_O_3_ raise it, contrary to expectations. We might assume that the temperature shift of T_iw_ can be related to changes in heat distribution in a sample with metal oxide powders, or in other words, to the technical features of the experiment. This hypothesis could be supported by the fact that ΔT_iw_ equals just a few hundredths of a degree. We checked it out by tracing the dependence of the T_iw_ temperature on different amount of CuO. The Figure 5 shows that T_iw_ did not depend on the amount of insoluble CuO powder in the test tube, despite the fact that its amount varied from 1.5 to 170 mg and the volume of this powder was quite large.

We cannot yet explain the increase of T_iw_ in the presence of some substances, however, we decided to discuss this effect, since in various researches a change in the temperature of T_iw_ by about 0.1°C is considered significant and is associated, for example, with the influence of dissolved antifreeze proteins (for example (Gupta & Deswal, 2014)).

Figure 5A also shows that the different amount of CuO in the test tube does not affect the temperature T_f_. This result is clear. One ice nucleator is enough for ice to form, and if there are several of them in a test tube (100 times more) then it will not affect the probability of ice nucleation for such a volume as 1ml (Finkelstein et al., 2022).

## Acknowledgements

We are grateful to Alexei V. Finkelstein for fruitful discussions.

This research was funded by Russian Science Foundation, grant number 21-14-00268

Authors declare no conflicts of interest.

## Notes

### Competing Interest Statement

The authors have declared no competing interest.

